# Agrochemical control of gene expression using evolved split RNA polymerase

**DOI:** 10.1101/2021.05.03.442273

**Authors:** Yuan Yuan, Jin Miao

**Affiliations:** Department of Neurophysiology and Neuropharmacology, Institute of Special Environmental Medicine and Co-innovation Center of Neuroregeneration, Nantong University, Nantong, 226019, China; Duke Kunshan University, 8 Duke Avenue, Kunshan, Jiangsu Province, 215316, China

## Abstract

Chemical inducible gene expression systems are valuable tools for rational control of gene expression both for basic research and biotechnology. However, most chemical inducers are confined to certain groups of organisms. Therefore, dissecting interaction between different organisms could be challenging using existing chemical inducible systems. We engineered a mandipropamid induced gene expression system (Mandi-T7) based on evolved split T7 RNAP system. As a proof-of-principle, we induced GFP expression in *E. coli* cells grown inside plant tissue.

## Introduction

Chemical inducible gene expression systems are powerful tools that allow conditional control over gene expression and facilitate study of toxic and essential genes. However, most chemical inducible systems were often designed to control gene expression in a single group of organisms in the laboratory condition. Working in complex context, like symbiosis between plant and bacteria, requires new chemical inducers that are neutral to different groups of organisms. Agrochemicals are attractive candidates because of their specific mode of action, low toxicity, and drug-like physical properties. Mandipropamid is the active ingredient of oomyceticide REVUS® and has been repurposed to control drought tolerance of plants through orthogonal control strategy.^1^ Mandipropamid and PYR1^MANDI^ protein constitute an engineered ligand-receptor pair, orthogonal to Abscisic acid (ABA) - PYR1 receptor pair.^1^ The Mandipropamid - PYR1^MANDI^ pair binds protein phosphatase HAB1 or ABI1 as the ABA-PYR1 pair (Figure S1).^1^ RNA polymerases which are modulated by protein-protein interaction would be desirable for coupling with Mandipropamid-sensing module to activate gene transcription. Split T7 RNA Polymerase (RNAP) has been evolved to be a proximity-dependent biosensor platform, which could transduce protein-protein interaction of light or small molecule sensing modules into activation of gene expression under the control of T7 promoter.^2^ ABA inducible CRISPR/Cas9 gene editing in mammalian cells has been demonstrated using this evolved split T7 RNAP system.^3^ Engineering Mandipropamid induced gene expression is feasible by modifying abscisic acid sensing module. Here, we report the repurposing of Mandipropamid to activate gene expression using the evolved split T7 RNA polymerase (Mandi-T7) and demonstrate the ability to regulate gene expression of bacteria grown inside plant tissue.

## Materials & Methods

### Plasmid construction

All genetic parts were synthesized and cloned into vectors by Genscript Biotechnology Inc. (Nanjing, China). Plasmid pET23a(+) was used to clone the Mandipropamid-T7 RNAP cassettes. And the backbone of compatible plasmid pCDFduet-1 was used to clone the T7pro-sfGFP reporter cassette. Sequence of all genetic parts were described in Table S2 and Table S3.

### Mandipropamid responsive assay for evolved split T7 RNA polymerase

Both driver and reporter plasmids were co-transformed into *E. coli* strain Top10 (TIANGEN biotech., Beijing, China). Single colonies were inoculated 2XYT medium with antibiotics at 37 °C. Overnight culture was transferred to fresh medium with antibiotics at 1:400 ratio. At the same time, Mandipropamid (sc-235565, Santa Cruz) or DMSO (solvent control) was added. The culture was incubated for 6 hours, and 100 µL of each sample was added to a 96 well plate and mixed with 190 µL 0.1% Tween-80. Both florescence (GFP, Ex: 488, Em: 510; mcherry, Ex: 587, Em: 630) and OD_600_ was measured by Perkin Elmer Enspire™ 2300 Multilabel Reader. After the values for medium were subtracted, florescence over OD_600_ was calculated and compared to that of the DMSO control samples.

### Mandipropamid responsive assay for split GFP

Plasmids were transformed into *E. coli* strain BL21(DE3) (TRANSGEN biotech., Beijing, China). Single colonies were inoculated 2XYT medium with antibiotics at 37 °C. Overnight culture was transferred to fresh medium at 1:100 ratio. When OD_600_ reached between 0.4 to 0.6, IPTG (1mM final) was added to induce expression of GFP1-9. Mandipropamid or DMSO (solvent control) was also added. The culture was incubated for 3 hours, and 100 µL of each sample was transferred to a 96 well plate. Both GFP florescence (Ex: 488, Em: 510) and OD_600_ was measured by Perkin Elmer Enspire™ 2300 Multilabel Reader. After the values for medium were subtracted, florescence over OD_600_ was calculated and compared to that of the DMSO control sample.

### Time course fluorescence measurement

*E. coli* Top10 strain containing Mandi-T7 and reporter plasmids was generated as mentioned above. Single colonies were inoculated 2XYT medium with antibiotics at 37 °C. Overnight culture was transferred to fresh medium with antibiotics at 1:200 ratio. When OD_600_ reached roughly 0.2, aliquots of 500 µL culture was distributed into individual wells of a deep-well plate or into the hollow septate stems of water spinach (*Ipomoea aquatica*). Mandipropamid (sc-235565, Santa Cruz) or DMSO (solvent control) was then added. The plate was incubated at 1000 rpm for 6 hours. Samples were harvested after 15 min, 30 min, 1 hour, 2 hour, 3 hour, 4 hour, and 6 hour. 100 µL of each sample was added to a 96 well plate. Both florescence (GFP: Ex: 488, Em: 510; mcherry: Ex: 587, Em: 630) and OD_600_ was measured by Perkin Elmer Enspire™ 2300 Multilabel Reader. After the values for medium were subtracted, florescence over OD_600_ was calculated.

### Western Blot analysis

*E*.*coli* culture was harvested at series of time points from deep-well plates using centrifuge. Samples were boiled in 1X loading buffer for 5 min. Relative protein concentration was determined by BCA Assay. Total protein was resolved using SDS-PAGE. The protein sample was transferred to PVDF membranes (Millipore) using Mini Trans-Blot (Bio-Rad). Anti-GFP (MF090, 1:2000 dilution) and anti-GAPDH (MF091, 1:1000 dilution) antibodies from Mei5bio were used.

## Results and discussion

To simplify circuit design and troubleshooting, we deployed the Mandi-T7 driver module and reporter module on two compatible plasmids. For the reporter module, expression of superfolder Green Fluorescent Protein (sfGFP) was driven by the T7 promoter. For the driver module, we followed the reported architecture of split T7 RNAP.^3^ Briefly, we fused PYR1^MANDI^ to the RNAP C-terminal half (T7 RNAPc) and the evolved RNAP N-terminal half (eRNAPn) to the N-terminal truncated ABI1 (Figure 1A). We used relatively long flexible linker (12 aa) to fuse PYR1^MANDI^ and T7 RNAPc for achieving high-level expression upon induction.^2^ We used the catalytic inactive mutant (D143A) of ABI1 protein (ABI) to minimize unintended effects.^4^ To avoid using additional inducer molecules, we used constitutive promoters to drive the expression of Mandi-T7. We chose standard promoter of medium strength (J23105) to drive the expression of PYR1^MANDI^-T7 RNAPc to avoid overwhelming the metabolism.^5^ To evaluate the performance of Mandi-T7 system, we measured the response of the Mandi-T7 to different concentration of Mandipropamid. The result showed a dose-dependent response, which saturated at 100 µM Mandipropamid (Figure 1B and Figure S2). A fold-induction of 44X was achieved, within the same order of magnitude as the 100X induction of the ABA-inducible RNAP sensor at 100 µM.^3^ The difference between the two systems is that ABA sensor can be further induced to 468X fold change at 10mM.^3^

**Figure 1.**
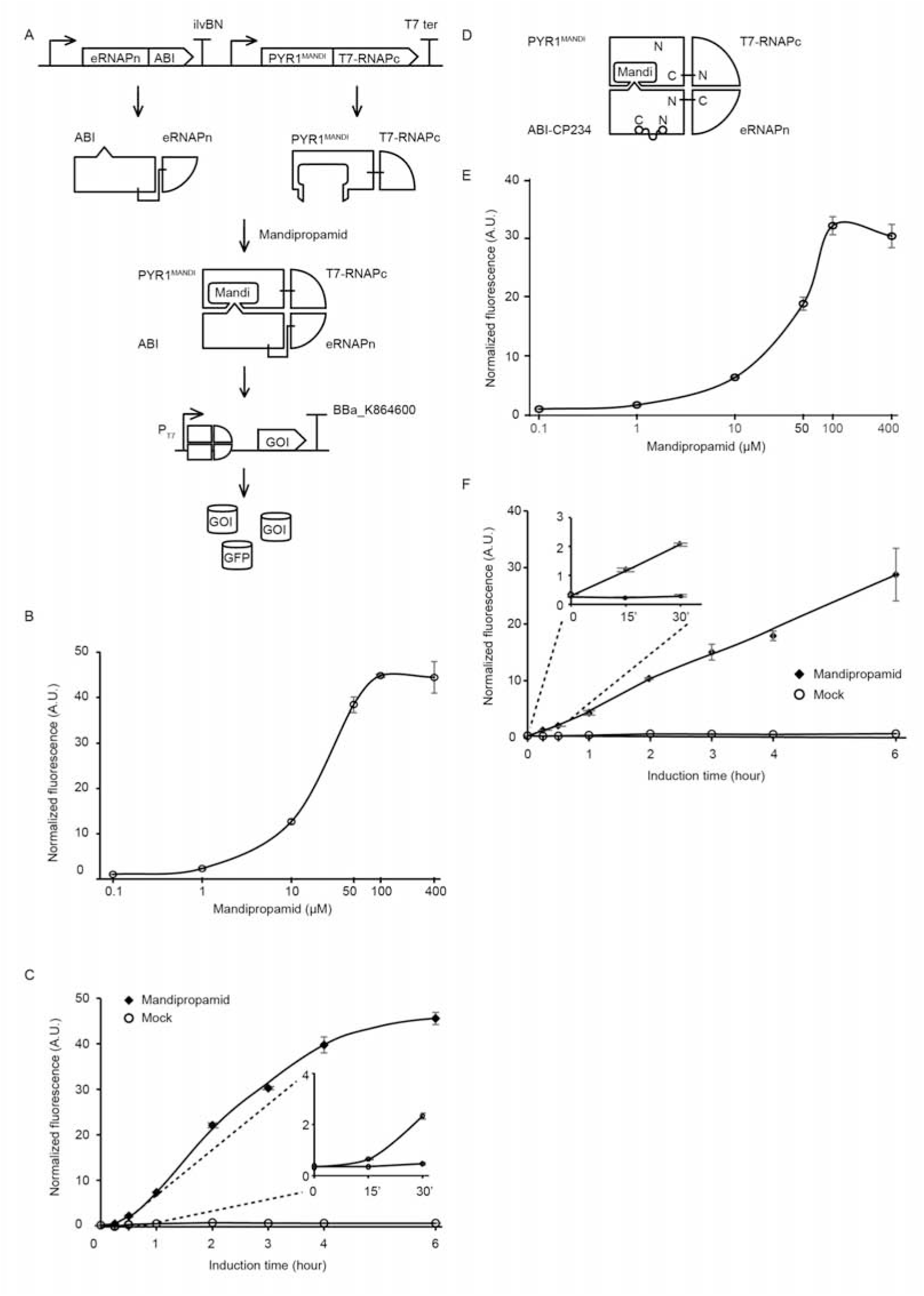
Engineering Mandi-T7 and its application. A, schematic of Mandi-T7 system and GFP reporter under control of T7 promoter. Bent arrow: promoter; Box arrow: CDS; Large T-shape: terminator. B, dose-response curve of Mandi-T7 in *E. coli* Top10. C, inducer response after 6 hours of induction in liquid culture. D, schematic of CP234-Mandi-T7. The original termini of ABI were linked by a flexible linker. E, dose-response curve of CP234-Mandi-T7 in *E. coli* Top10. F, time course after induction for 6 hours in the hollow septate stems of water spinach.

To understand kinetic characteristics of Mandi-T7 system, we measured fluorescence intensity over the course of induction for 6 hours. Fluorescence above the background level was detected at 15 min (Figure 1C). The induction plateaued after 6 hours (Figure 1C). We also detected fluorescence signal without induction (Figure 1C), which may be caused by the residual background activity of the evolved split-T7 polymerase.^3^ We also tested mcherry as the reporter. The dose-response curve of mcherry showed 140X induction at 100 µM (Figure S3A). The time course expression analysis showed induction plateaued after 4 hours (Figure S3B). The increased fold change for mcherry is consistent with low cellular autofluorescence at its emission region.^6^ We failed to detect toxic effect of Mandipropamid on *E*.*coli* growth (Figure S4).

In a separate attempt, we tried a circular permutated version of ABI protein, which provides alternative topology for complementation of the split T7 RNAP. We noticed that Ser 234 at the end of helix 2 is closer to the C terminus of PYR1^MANDI^ than the original N-terminus (Figure S5). We joined the original N and C termini of ABI by a flexible linker and generated the circular permutant CP234, for which Ser234 of ABI serves as the new N-terminus (Figure 1D). To test the retention of the activity to bind PYR1^MANDI^, we used the tripartite split-GFP based system because of its low background fluorescence and minimal effect on fusion protein.^7^ We observed concentration dependent increase in GFP florescence after Mandipropamid induction (Figure S6). This result indicated that ABI-CP234 retained the ability to interact with PYR1^MANDI^ in the presence of Mandipropamid. Next, we adapted ABI-CP234 to the evolved split T7 RNAP system (CP234-Mandi-T7), replacing ABI using ABI-CP234. The result also showed strong sfGFP expression upon Mandipropamid induction like the Mandi-T7 system, with a fold induction of 33 (Figure 1E and Figure S2).

To provide a proof-of-principle that Mandi-T7 is useful in complex setting, we evaluated the ability of Mandi-T7 to induce bacterial gene expression inside the hollow septate stems of water spinach (*Ipomoea aquatica*). We followed reporter expression using fluorescence intensity and Western Blot. The result showed induction after 15 min and induction level kept on increasing over 6-hour period (Figure 1F, Figure S7). This result suggested that Mandi-T7 has the potential to enable manipulation of gene expression of plant-associated microorganisms.

In summary, this study provides a new tool for agrochemical control gene expression. We showed that Mandipropamid inducible gene expression can be readily built based on the evolved split T7 RNAP system. T7 RNAP is orthogonal and widely applicable to many prokaryotic and eukaryotic organisms. Mandi-T7 can be further integrated with other tools like CRISPRi technology. Low toxicity, easy absorption, and cost-effectiveness of Mandipropamid will give Mandi-T7 leverage in other complex contexts like vector-borne bacterial plant pathogens. We also tested alternative topology (ABI-CP234) for generating ABI fusion protein and hope this result might help expand Mandipropamid to other proximity-dependent systems. CP234-ABI might also be applicable to ABA FRET sensor. The reporter signal in uninduced conditions could be caused by constitutive expression of Mani-T7 module and low level self-assembly of split T7 RNAP. Further engineering efforts like directed evolution are needed to lower the basal expression level and increase the dynamic range. We hope the Mandi-T7 system will expand rational control over gene expression to diverse biological context.

## Acknowledgements

We would like to thank Dr. Jinyue Pu and Professor Bryan Dickinson (University of Chicago) for sharing plasmids and vector information. We want to thank Dr. Xin Zhou (Nantong University) for technical help with Confocal imaging. We also want to thank Professor Linfeng Huang (Duke Kunshan University) for advices 163 and generous sharing equipment and reagents. This work was partly supported by the 164 National Natural Science Foundation of China (Grant 31500690 to J. M.).

