## Supplemental Data for "Agrochemical control of gene expression using evolved split RNA polymerase"

for

Summary

This PDF file includes:

Supplementary Figure S1 to S7

Supplementary Table S1 and S3

**Supplementary figures**

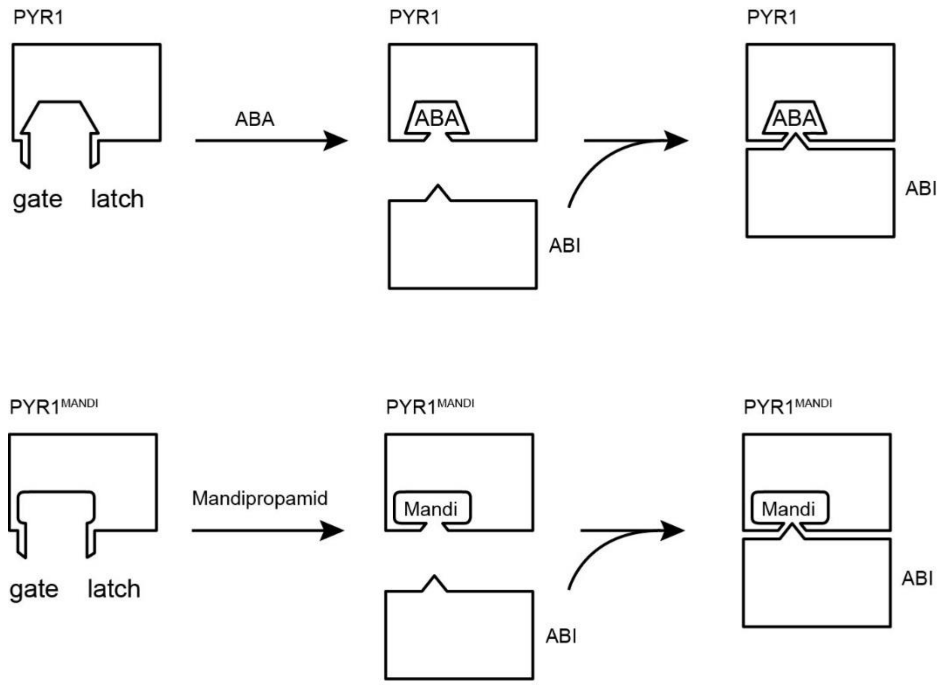


Figure S1, Mandipropamid - PYR1$^{MANDI}$ is an engineered orthogonal ligand receptor pair of ABA - PYR1. Mechanism of mandipropamid perception (based on Park, 2015) is similar to the gate-latch-lock mechanism of ABA preception (Melcher, 2009). Binding of Mandipropamid molecule leads to conformational change of PYR1$^{MANDI}$, like the closure of gate and latch. The surface created after the closure enables competitive binding to the active site of ABI (W300), which is depicted as a protrusion.

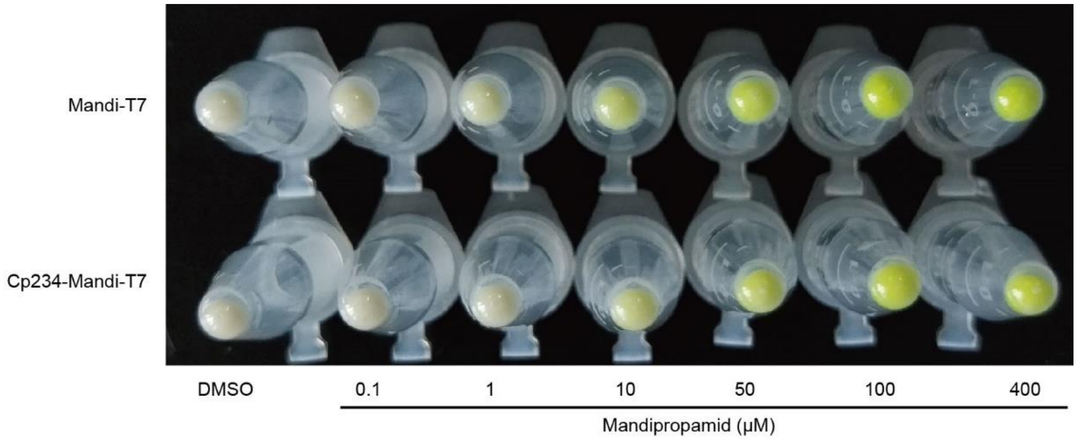


Figure S2, cell pellet of *E. coli* culture 6 hr post-induction. Picture was taken under normal day light condition.

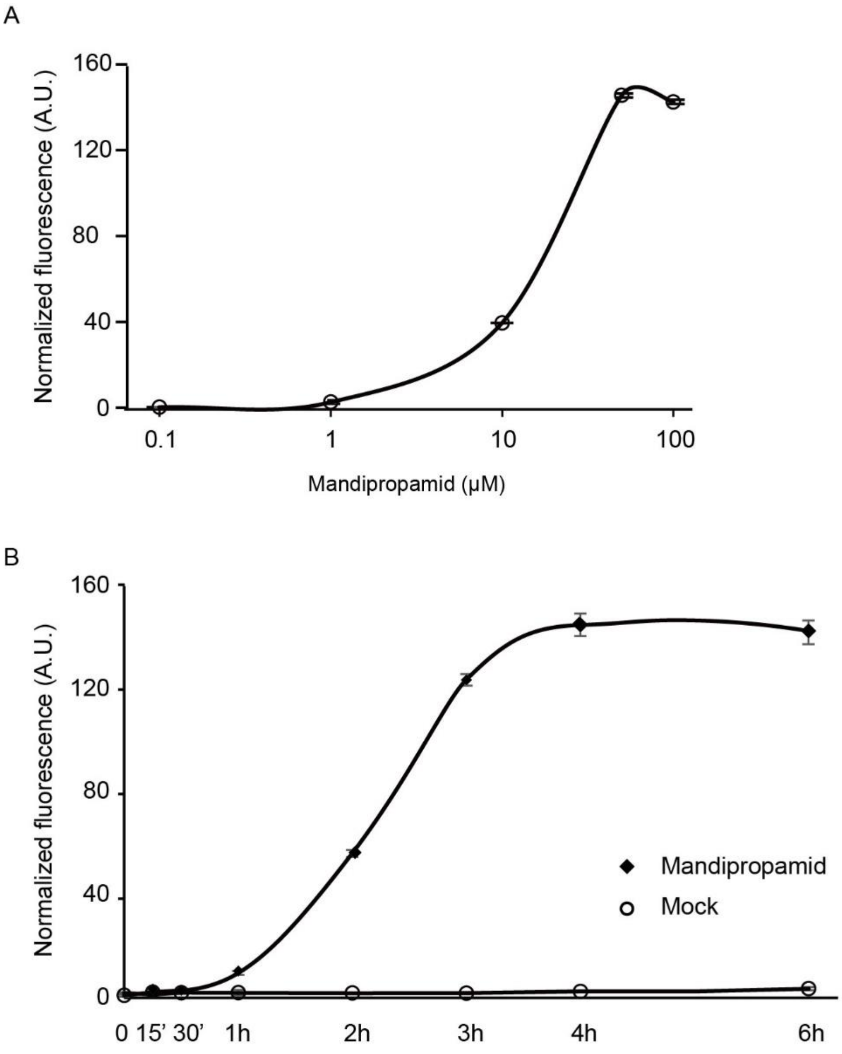


Figure S3, dose response and time course analysis using mcherry as the reporter. A, induction of mcherry expression with different concentrations of Mandipropamid. B, expression of mcherry over the time course of experiment after inducer or mock was added.

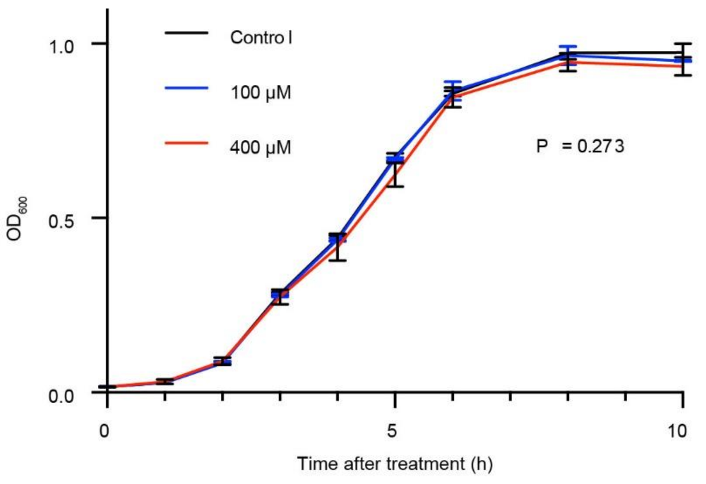


Figure S4, growth curve of *E. coli* treated with Mandipropamid (100 μM and 400 μM). Data was analyzed using a two-way repeated measures ANOVA.

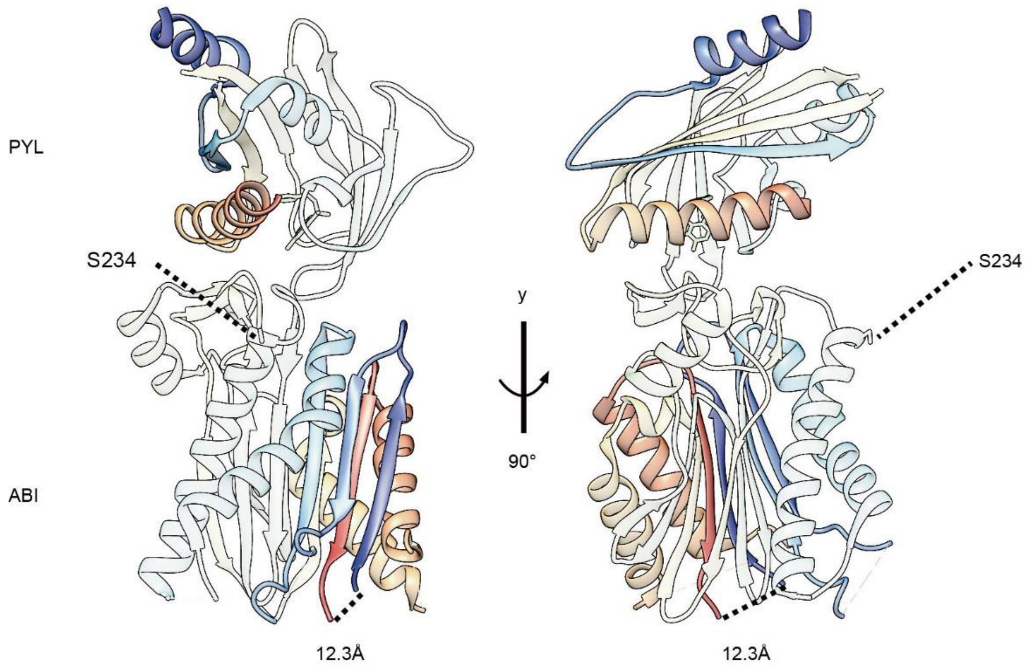


Figure S5, Ser234 of ABI is closer to the C-terminus of PYL than the N terminus. Peptide chains are shown in rainbow color from blue at the N terminus to red at the C terminus. The distance between original termini of ABI is 12.3Å. The cartoon was generated based on the structure of ABI-PYL-Pyrabactin complex (PDB ID: 3nmn) using Chimera 1.15.

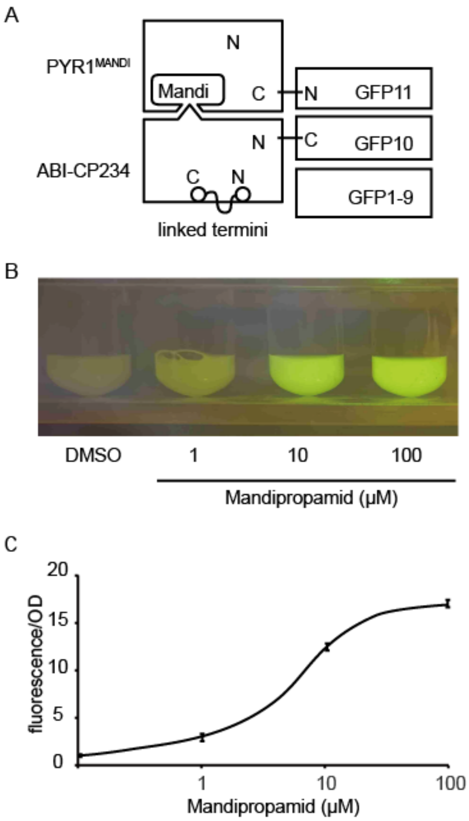


Figure S6, The tripartite split-GFP assay for ABI-CP234. A, Schematic of the tripartite split-GFP assay. The original termini of ABI were linked by a flexible linker. GFP protein is reconstituted only if GFP10 and GFP 11 fragments are brought into close proximity. B, photo of *E. coli* liquid culture taken with orange filter illuminated by blue light. C, dose-response curve of the tripartite split-GFP assay.

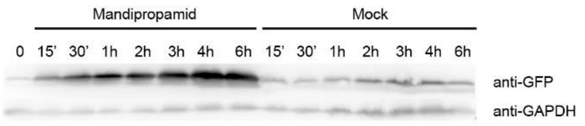


Figure S7, Western blot of GFP reporter expression over the time course of the induction experiment.

Table S1: comparison of the T7-eRNAP based ABA biosensor system (ABA for simplicity) with Mandi and CP234-Mandi.

| system | module | expression cassette | promoter | marker | origin |
|---|---|---|---|---|---|
| ABA (Pu *et al.*, 2018) | evolved T7N-ABI | 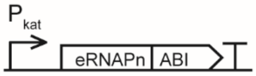  | constitutive | Spec | P15A |
| | PYR1-T7C | 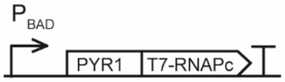  | inducible | Chlr | CloDF13 |
| | luciferase reporter | 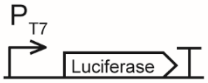  | | Carb | SC101 |
| Mandi-T7 | evolved T7N-ABI; PYR1$^{MANDI}$-T7C | 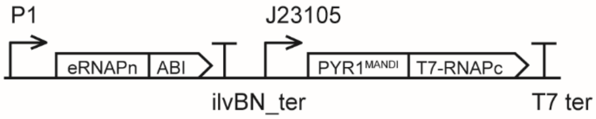  | constitutive | Amp | pBR322 |
| | sfGFP reporter | 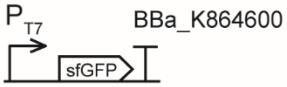  | | Spec | CloDF13 |
| CP234-Mandi-T7 | evolved T7N-ABI; PYR1$^{MANDI}$-T7C | 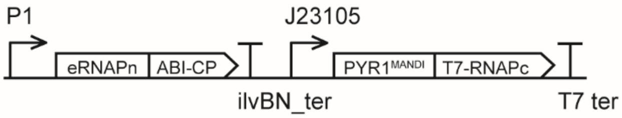  | constitutive | Amp | pBR322 |
| | sfGFP reporter | 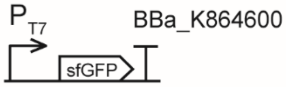  | | Spec | CloDF13 |

Table S2: sequence and source of genetic parts

| Name | Sequence | Source |
|---|---|---|
| P1 | tgttcacattcgaaccgtctctgctttgacatcttatgattctcgactgtaaagtcgtggcca | 1 |
| P2 | tgttcacattcgaaccgtctctgctttgacaacatgctgtgcggtgttgtaaagtcgtggccaggagaatacgacag | 1 |
| RBS1 | caacgctgcacccgaatcacattacggactattatt | 2 |
| RBS2 | gcaattgcaagaaggaggatattg | 2 |
| T7-eRNAPN (d5-19) | MNTINIAKNDFSDIELAAIPLNTLADHYGERSARGQLALEHESYEMGEARFRKMFECQLKAGKVAD NAAAKPLITTLLPKMIARINDWFEEVKAKRGRRPTAFKFLKEIKPEAVAYITIKTSLACLTSADNTTVQ AVASAIGRTIEDEARFGRIRDLEAKHFKKNVEEQLNKRVGHVYK | 3 |
| ABI | VPLYGFTSICGRRPEMEAAVSTIPRFLQSSSGSMLDGRFDPQSAAHFFGVYDGHGGSQVANYCRE RMHLALAEEIAKEKPMLCDGDTWLEKWKKALFNSFLRVDSEIESVAPETVGSTSVVAVVFPSHIFVA NCGDSRAVLCRGKTALPLSVDHKPDREDEAARIEAAGGKVIQWNGARVFGVLAMSRSIGDRYLKP SIIPDPEVTAVKRVKEDDCLILASDGVWDVMTDEEACEMARKRILLWHKKNAVAGDASLLADERRK EGKDPAAMSAAEYLSKLAIQRGSKDNISVVVVDLK (AT4G26080.1, V126-K423, D143A) | 3, 4 |
| ABI-CP234 (circularly permutated) | SVAPETVGSTSVVAVVFPSHIFVANCGDSRAVLCRGKTALPLSVDHKPDREDEAARIEAAGGKVIQ WNGARVFGVLAMSRSIGDRYLKPSIIPDPEVTAVKRVKEDDCLILASDGVWDVMTDEEACEMARK RILLWHKKNAVAGDASLLADERRKEGKDPAAMSAAEYLSKLAIQRGSKDNISVVVVDLKGGSGSGS SVPLYGFTSICGRRPEMEAAVSTIPRFLQSSSGSMLDGRFDPQSAAHFFGVYDGHGGSQVANYCR ERMHLALAEEIAKEKPMLCDGDTWLEKWKKALFNSFLRVDSEIE (S234 -K423-linker-V126-E233, D143A) | 3 |
| Terminator 1 | attcaagacccccgcaccgaaaggtccgggggtttttttacta | 5 |
| PYR1[MANDI] | MPSELTPEERSELKNSIAEFHTYQLDPGSCSSLHAQRIHAPPELVWSIVRRFDKPQTHRHFIKSCSV EQNFEMRVGCTRDIIVISGLPANTSTERLDILDDERRVTGASIIGGEHRLTNYKGVTTVHRFEKENRI WTVVLESYVVDMPEGNSEDDTRMLADTVVKLNLQKLATVAEAMA | 6 |
| T7-RNAPC | KAFMQVVEADMLSKGLLGGEAWSSWHKEDSIHVGVRCIEMLIESTGMVSLHRQNAGVVGQDSETI ELAPEYAEAIATRAGALAGISPMFQPCVVPPKPWTGITGGGYWANGRRPLALVRTHSKKALMRYE DVYMPEVYKAINIAQNTAWKINKKVLAVANVITKWKHCPVEDIPAIEREELPMKPEDIDMNPEALTA WKRAAAAVYRKDKARKSRRISLEFMLEQANKFANHKAIWFPYNMDWRGRVYAVSMFNPQGNDM TKGLLTLAKGKPIGKEGYYWLKIHGANCAGVDKVPFPERIKFIEENHENIMACAKSPLENTWWAEQ DSPFCFLAFCFEYAGVQHHGLSYNCSLPLAFDGSCSGIQHFSAMLRDEVGGRAVNLLPSETVQDIY GIVAKKVNEILQADAINGTDNEVVTVTDENTGEISEKVKLGTKALAGQWLAYGVTRSVTKRSVMTLA YGSKEFGFRQQVLEDTIQPAIDSGKGLMFTQPNQAAGYMAKLIWESVSVTVVAAVEAMNWLKSAA KLLAAEVKDKKTGEILRKRCAVHWVTPDGFPVWQEYKKPIQTRLNLMFLGQFRLQPTINTNKDSEID AHKQESGIAPNFVHSQDGSHLRKTVVWAHEKYGIESFALIHDSFGTIPADAANLFKAVRETMVDTYE SCDVLADFYDQFADQLHESQLDKMPALPAKGNLNLRDILESDFAFA* | 3 |
| Terminator 2 | ctagcataaccccttggggcctctaaacgggtcttgaggggttttttg | pET23a |
| GFP (sfGFP) | MRKGEELFTGVVPILVELDGDVNGHKFSVRGEGEGDATNGKLTLKFICTTGKLPVPWPTLVTTLTY GVQCFARYPDHMKQHDFFKSAMPEGYVQERTISFKDDGTYKTRAEVKFEGDTLVNRIELKGIDFKE DGNILGHKLEYNFNSHNVYITADKQKNGIKANFKIRHNVEDGSVQLADHYQQNTPIGDGPVLLPDN HYLSTQSVLSKDPNEKRDHMVLLEFVTAAGITHGMDELYK | 1 |
| T7 pro_RBS (for sfGFP) | taatacgactcactatagggagaccacaacggtttccctctaCaaataattttgtttaactttaagaaggagatatacat | pET23a |
| GFP10 | MDLPDDHYLSTQTILSKDLN | 7 |
| GFP11 | EKRDHMVLLEYVTAAGITDAS | 7 |
| GFP1-9 | MRKGEELFTGIVPILVELDGDVNGHKFFVRGEGEGDATIGKLSLKFICTTGKLPVPWPTLVTTLTYG VQCFSRYPDHMKRHDFFKSAMPEGYVQERTIYFKDDGTYKTRAEVKFEGDTLVNRIELKGIDFKED GNILGHKLEYNFNSHKVYITADKQNNGIKANFTIRHNVEDGSVQLADHYQQNTPIGDGPVLLP | 7 |
| T7_lacO_RBS (for GFP1-9) | taatacgactcactataggggaattgtgagcggataacaattcccctctagaaataattttgtttaactttaagaaggagatatacc | pET28a |
| J23109 | tttacagctagctcagtcctagggactgtgctagct | Biobrick |
| J23105 | tttacggctagctcagtcctaggtactatgctagct | Biobrick |
| mcherry | MVSKGEEDNMAIIKEFMRFKVHMEGSVNGHEFEIEGEGEGRPYEGTQTAKLKVTKGGPLPFAWDI LSPQFMYGSKAYVKHPADIPDYLKLSFPEGFKWERVMNFEDGGVVTVTQDSSLQDGEFIYKVKLR GTNFPSDGPVMQKKTMGWEASSERMYPEDGALKGEIKQRLKLKDGGHYDAEVKTTYKAKKPVQL PGAYNVNIKLDITSHNEDYTIVEQYERAEGRHSTGGMDELYK | |

Table S3: sequence of plasmids

| | |
|---|---|
| pCDF-T7-sfGFP<br>(Turquoise:<br>T7 promoter<br>yellow:<br>sfGFP<br>purple:<br>Terminator) | gcgaaattaatacgactcactatagggagaccacaacggtttccctctaCaaataattttgtttaactttaagaaggagatatacatatgcgtaaaggcg<br>aagagctgttcactggtgtcgtccctattctggtggaactggatggtgatgtcaacggtcataagttttccgtgcgtggcgagggtgaaggtgacgcaact<br>aatggtaaactgacgctgaagttcatctgtactactggtaaactgccagtaccttggccgactctggtaacgacgctgacttatggtgttcagtgctttgctc<br>gttatccggaccacatgaagcagcatgacttcttcaagtccgccatgccggaaggctatgtgcaggaacgcacgatttcatttaaggatgacggcacgt<br>acaaaacgcgtgcggaagtgaaatttgaaggcgataccctggtaaaccgcattgagctgaaaggcattgactttaaagaagacggcaatatcctgg<br>gccataagctggaatacaattttaacagccacaatgtttacatcaccgccgataaacaaaaaaatggcattaaagcgaatttcaaaattcgccacaac<br>gtggaggatggcagcgtgcagctggctgatcactaccagcaaaacactccaatcggtgatggtcctgttctgctgccagacaatcactatctgagcac<br>gcaaagcgttctgtctaaagatccgaacgagaaacgcgatcacatggttctgctggagttcgtaaccgcagcgggcatcacgcacggtatggatgaa<br>ctgtacaaatgagaattcttgttcagaacgctcggtcttgcacaccgggcgttttttctttgtgagtccaatccaaaacgccgcgttcagcggcgtttttctgc<br>ttctgaaag |
| pJM135A<br>(Turquoise:<br>P1<br>Yellow:<br>eT7N-ABI<br>red:<br>Terminator 1<br>Teal:<br>J23105<br>Green:<br>PYR1$^{MANDI}$-T7C<br>Blue:<br>Terminator 2<br>) | tgttcacattcgaaccgtctctgctttgacatcttatgattctcgactgtaaagtcgtggccacaacgctgcacccgaatcacattacggactattatatgaa<br>cacgattaacatcgctaagaacgacttctctgacatcgaactggctgctatcccgctcaacactctggctgaccattacggtgagcgttcagctcgcgga<br>cagttggcccttgagcatgagtcttacgagatgggtgaagcacgcttccgcaagatgtttgagtgtcaacttaaagctggtaaggttgcggataacgctg<br>ccgccaagcctctcatcactaccctcctccctaagatgattgcccgcatcaacgactggtttgaggaagtgaaagctaagcgcggcaggcgcccgac<br>agccttcaagttcctcaaagaaatcaagccggaagccgtagcgtacatcaccattaagacctctctggcttgcctcaccagtgctgacaatacaaccgt<br>tcaggctgtagcaagcgcaatcggtcggaccattgaggacgaggctcgcttcggtcgtatccgtgaccttgaagctaagcacttcaagaaaaacgttg<br>aggaacaactcaacaagcgcgtagggcacgtctacaagggtggatccggctctggttcgagtgtcccccctgtatgggttcaccagcatttgtgggcgg<br>agaccagagatggaggcagccgtcagcactattccccggttcctgcagtcaagctccggcagcatgctggacgggaggttcgatccacagtccgcc<br>gctcacttctttggggtctacgatggacatggcgggtcccaggtggccaactattgcagggagcgcatgcacctggctctggcagaggaaatcgccaa<br>ggagaaacctatgctgtgcgacggagatacatggctggaaaagtggaagaaagccctgttcaactcttttctgcgggtggactccgagatcgaatctgt<br>cgctcctgagaccgtgggctctacaagtgtggtcgcagtggtcttcccatctcacatttttgtcgcaaattgcggcgacagccgggccgtgctgtgcaggg<br>gaaagaccgccctgccactgagcgtggaccataaacccgacagggaggatgaagcagcccgcatcgaggctgcaggaggcaaagtgatccagt<br>ggaacggagcacgggtgtttggcgtcctggccatgtcacgcagcattggggatcgatacctgaaaccatcaatcattcccgaccctgaagtgactgcc<br>gtcaagagagtgaaagaagacgattgcctgatcctggctagcgacggcgtctgggatgtgatgaccgacgaggaagcatgtgagatggcccgaaa<br>gcggattctgctgtggcataagaaaaatgccgtggctggcgatgctagtctgctggcagacgagcggagaaaggaagggaaagatcccgccgctat<br>gagtgcagccgaatatctgtcaaaactggctattcagaggggcagtaaggacaacatctccgtcgtcgtcgtggacctgaagtgaggtaccattcaag<br>acccccgcaccgaaaggtccggggtttttttttactatttaaatcctgccagtttacggctagctcagtcctaggtactatgctagctgcaattgcaagaag<br>gaggatattgatgccatcggaacttacaccggaggaaaggtccgagttgaagaacagcattgcggagtttcatacatatcaactcgatccggggagtt<br>gttctagtttgcacgcccaaaggatacacgccccaccggaacttgtgtggagtatcgtccgccggttcgataaaccacaaacacacaggcatttcatta<br>aaagctgctccgtggagcagaactttgaaatgagagtgggatgcacacgggatataatagttataagcggcctcccggccaacacgagcaccgaa<br>agattggacatactggatgacgagcgccgcgtgacgggggcatcaataatcggtggggaacatcgccttacgaattataaaggcgtcactactgttca<br>tagatttgagaaggaaaatagaatctggactgttgttctggaatcttatgtggttgatatgccagagggtaactcggaagatgatacacgcatgctggcg<br>gacacggttgtgaaacttaatctgcaaaagctggcaactgtcgccgaagccatggcatcaggatctggtagcgcaggatcaggctcgagtggtaaag<br>catttatgcaagttgtcgaggctgacatgctctctaagggtctcctcggtggcgaggcgtggtcttcgtggcataaggaagactctattcatgtaggagtac<br>gctgcatcgagatgctcattgagtcaaccggaatggttagcctccaccgccaaaatgctggcgtagtaggtcaagactctgagactatcgaactcgca<br>cctgaatacgctgaggctatcgcaacccgtgcaggtgcgctggctggcatctctccgatgttccaaccttgcgtagttcctcctaagccgtggactggcat<br>tactggtggtggctattgggctaacggtcgtcgtcctctggcgctggtgcgtactcacagtaagaaagcactgatgcgctacgaagacgtttacatgcctg<br>aggtgtacaaagcgattaacattgcgcaaaacaccgcatggaaaatcaacaagaaagtcctcgcggtcgccaacgtaatcaccaagtggaagcat<br>tgtccggtcgaggacatccctgcgattgagcgtgaagaactcccgatgaaaccggaagacatcgacatgaatcctgaggctctcaccgcgtggaaa<br>cgtgctgccgctgctgtgtaccgcaaggacaaggctcgcaagtctcgccgtatcagccttgagttcatgcttgagcaagccaataagtttgctaaccata<br>aggccatctggttcccttacaacatggactggcgcggtcgtgtttacgctgtgtcaatgttcaacccgcaaggtaacgatatgaccaaaggactgcttac<br>gctggcgaaaggtaaaccaatcggtaaggaaggttactactggctgaaaatccacggtgcaaactgtgcgggtgtcgataaggttccgttccctgagc<br>gcatcaagttcattgaggaaaaccacgagaacatcatggcttgcgctaagtctccactggagaacacttggtgggctgagcaagattctccgttctgctt<br>ccttgcgttctgctttgagtacgctggggtacagcaccacggcctgagctataactgctcccttccgctggcgtttgacgggtcttgctctggcatccagca<br>cttctccgcgatgctccgagatgaggtaggtggtcgcgcggttaacttgcttcctagtgaaaccgttcaggacatctacgggattgttgctaagaaagtca<br>acgagattctccaagcagacgcaatcaatgggaccgataacgaagtagttaccgtgaccgatgagaacactggtgaaatctctgagaaagtcaagc<br>tgggcactaaggcactggctggtcaatggctggcttacggtgttactcgcagtgtgactaagcgttcagtcatgacgctggcttacgggtccaaagagtt<br>cggcttccgtcaacaagtgctggaagataccattcagccagctattgattccggcaagggtctgatgttcactcagccgaatcaggctgctggatacatg<br>gctaagctgatttgggaatctgtgagcgtgacggtggtagctgcggttgaagcaatgaactggcttaagtctgctgctaagctgctggctgctgaggtca<br>aagataagaagactggagagattcttcgcaagcgttgcgctgtgcattgggtaactcctgatggtttccctgtgtggcaggaatacaagaagcctattca<br>gacgcgcttgaacctgatgttcctcggtcagttccgcctccagcctaccattaacaccaacaaagatagcgagattgatgcacacaaacaggagtctg<br>gtatcgctcctaactttgtacacagccaagacggtagccaccttcgtaagactgtagtgtgggcacacgagaagtacggaatcgaatcttttgcactgatt<br>cacgactccttcggtaccattccggctgacgctgcgaacctgttcaaagcagtgcgcgaaactatggttgacacatatgagtcttgtgatgtactggctga<br>tttctacgaccagttcgctgaccagttgcacgagtctcaattggacaaaatgccagcacttccggctaaaggtaacttgaacctccgtgacatcctcgagt<br>cggacttcgcgttcgcgtaatagcaccaccaccaccaccactgagatccggctgctaacaaagcccgaaaggaagctgagttggctgctgccaccg<br>ctgagcaataactagcataaccccttggggcctctaaacgggtcttgaggggttttttg |
| pCDF-T7-lacO-<br>GFP1-9<br>(Turquoise:<br>T7 promoter<br>Red:<br>lacO<br>yellow:<br>GFP1-9<br>Violet:<br>terminator) | TAATACGACTCACTATAGGGGAATTGTGAGCGGATAACAATTCCCCTCTAGAAATAATTTTGTTTAACTT<br>TAAGAAGGAGATATACCatgcgcaaaggcgaagaactgtttaccggcattgtgccgattctggtggaactggatggcgatgtgaacggcca<br>taaatttttttgtgcgcggcgaaggcgaaggcgatgcgaccattggcaaactgagcctgaaatttatttgcaccaccggcaaactgccggtgccgtggcc<br>gaccctggtgaccaccctgacctatggcgtgcagtgctttagccgctatccggatcatatgaaacgccatgatttttttaaaagcgcgatgccggaaggc<br>tatgtgcaggaacgcaccatttattttaaagatgatggcacctataaaacccgcgcggaagtgaaatttgaaggcgataccctggtgaaccgcattgaa<br>ctgaaaggcattgattttaaagaagatggcaacattctgggccataaactggaatataactttaacagccataaagtgtatattaccgcggataaacag<br>aacaacggcattaaagcgaactttaccattcgccataacgtggaagatggcagcgtgcagctggcggatcattatcagcagaacaccccgattggcg<br>atggcccggttcttcttccttaggaattcttgttcagaacgctcggtcttgcacaccgggcgttttttctttgtgagtcca |
| pJM134<br>(Turquoise:<br>J23101 | tttacagctagctcagtcctaggtattatgctagcgaaattacccttgccttaactaataagagagctgtctatggacctgcctgacgaccactacctgtcc<br>acccagaccatcctgtccaaggacctgaacggatctggtggatccggctctggttcgagttctgtcgctcctgagaccgtgggctctacaagtgtggtcg<br>cagtggtcttcccatctcacatttttgtcgcaaattgcggcgacagccgggccgtgctgtgcaggggaaagaccgccctgccactgagcgtggaccata |

| | |
|---|---|
| Yellow:<br>G10- ABI-CP1<br>Red:<br>Terminator 1<br>Violet:<br>P2<br>Green:<br>PYR1$^{MANDI}$-G11<br>Gray:<br>Terminator 2) | aacccgacagggaggatgaagcagcccgcatcgaggctgcaggaggcaaagtgatccagtggaacggagcacgggtgtttggcgtcctggccat<br>gtcacgcagcattggggatcgatacctgaaaccatcaatcattcccgaccctgaagtgactgccgtcaagagagtgaaagaagacgattgcctgatc<br>ctggctagcgacggcgtctgggatgtgatgaccgacgaggaagcatgtgagatggcccgaaagcggattctgctgtggcataagaaaaatgccgtg<br>gctggcgatgctagtctgctggcagacgagcggagaaaggaagggaaagatcccgccgctatgagtgcagccgaatatctgtcaaaactggctatt<br>cagaggggcagtaaggacaacatctccgtcgtcgtcgtggacctgaagggcggtagtggttcgggttcttcggtcccccctgtatgggttcaccagcattt<br>gtgggcggagaccagagatggaggcagccgtcagcactattccccggttcctgcagtcaagctccggcagcatgctggacgggaggttcgatccac<br>agtccgccgctcacttctttggggtctacgatggacatggcgggtcccaggtggccaactattgcagggagcgcatgcacctggctctggcagaggaa<br>atcgccaaggagaaacctatgctgtgcgacggagatacatggctggaaaagtggaagaaagccctgttcaactcttttctgcgggtggactccgagat<br>cgaatgaggtaccattcaagacccccgcaccgaaaggtccggggggttttttttactatttaaatcctgccagtgttcacattcgaaccgtctctgctttgaca<br>acatgctgtgcggtgttgtaaagtcgtggccaggagaatacgacaggcaattgcaagaaggaggatattgatgccatcggaacttacaccggagga<br>aaggtccgagttgaagaacagcattgcggagtttcatacatatcaactcgatccggggagttgttctagtttgcacgcccaaaggatacacgccccacc<br>ggaacttgtgtggagtatcgtccgccggttcgataaaccacaaacacacaggcatttcattaaaagctgctccgtggagcagaactttgaaatgagagt<br>gggatgcacacgggatataatagttataagcggcctcccggccaacacgagcaccgaaagattggacatactggatgacgagcgccgcgtgacg<br>gggggcatcaataatcggtggggaacatcgccttacgaattataaaggcgtcactactgttcatagatttgagaaggaaaatagaatctggactgttgttct<br>ggaatcttatgtggttgatatgccagagggtaactcggaagatgatacacgcatgctggcggacacggttgtgaaacttaatctgcaaaagctggcaa<br>ctgtcgccgaagccatggcaggatcaggctcgagtggttcaggatctggtgcaagcgaaaagcgagaccatatggttttgcttgagtatgttacagcgg<br>ctggcattaccgatgcatcatagtcggacttcgcgttcgcgtaatagcaccaccaccaccaccactgagatccggctgctaacaaagcccgaaagga<br>agctgagttggctgctgccaccgctgagcaataactagcataaccccttggggcctctaaacgggtcttgaggggttttttg |
| pJM1B6<br>(turquoise:<br>P1<br>Yellow:<br>eT7N-<br>ABI_CP234<br>Red:<br>Terminator 1<br>Teal:<br>P2<br>Green:<br>PYR1$^{MANDI}$-T7C<br>Blue:<br>Terminator 2<br>) | ctatactcggatatctgttcacattcgaaccgtctctgctttgacatcttatgattctcgactgtaaagtcgtggccacaacgctgcacccgaatcacattacg<br>gactattattatgaacacgattaacatcgctaagaacgacttctctgacatcgaactggctgctatcccgctcaacactctggctgaccattacggtgagc<br>gttcagctcgcggacagttggcccttgagcatgagtcttacgagatgggtgaagcacgcttccgcaagatgtttgagtgtcaacttaaagctggtaaggtt<br>gcggataacgctgccgccaagcctctcatcactaccctcctccctaagatgattgcccgcatcaacgactggtttgaggaagtgaaagctaagcgcgg<br>caggcgcccgacagccttcaagttcctcaaagaaatcaagccggaagccgtagcgtacatcaccattaagacctctctggcttgcctcaccagtgctg<br>acaatacaaccgttcaggctgtagcaagcgcaatcggtcggaccattgaggacgaggctcgcttcggtcgtatccgtgaccttgaagctaagcacttc<br>aagaaaaacgttgaggaacaactcaacaagcgcgtagggcacgtctacaagggtggatccggctctggttcgagttctgtcgctcctgagaccgtgg<br>gctctacaagtgtggtcgcagtggtcttcccatctcacatttttgtcgcaaattgcggcgacagccgggccgtgctgtgcaggggaaagaccgccctgcc<br>actgagcgtggaccataaacccgacagggaggatgaagcagcccgcatcgaggctgcaggaggcaaagtgatccagtggaacggagcacggg<br>tgtttggcgtcctggccatgtcacgcagcattggggatcgatacctgaaaccatcaatcattcccgaccctgaagtgactgccgtcaagagagtgaaag<br>aagacgattgcctgatcctggctagcgacggcgtctgggatgtgatgaccgacgaggaagcatgtgagatggcccgaaagcggattctgctgtggca<br>taagaaaaatgccgtggctggcgatgctagtctgctggcagacgagcggagaaaggaagggaaagatcccgccgctatgagtgcagccgaatatc<br>tgtcaaaactggctattcagaggggcagtaaggacaacatctccgtcgtcgtcgtggacctgaagggcggtagtggttcgggttcttcggtcccccctgtat<br>gggttcaccagcatttgtgggcggagaccagagatggaggcagccgtcagcactattccccggttcctgcagtcaagctccggcagcatgctggacg<br>ggaggttcgatccacagtccgccgctcacttctttggggtctacgatggacatggcgggtcccaggtggccaactattgcagggagcgcatgcacctg<br>gctctggcagaggaaatcgccaaggagaaacctatgctgtgcgacggagatacatggctggaaaagtggaagaaagccctgttcaactcttttctgc<br>gggtggactccgagatcgaatgaggtaccattcaagacccccgcaccgaaaggtccggggggttttttttactatttaaatcctgccagtgttcacattcga<br>accgtctctgctttgacaacatgctgtgcggtgttgtaaagtcgtggccaggagaatacgacaggcaattgcaagaaggaggatattgatgccatcgg<br>aacttacaccggaggaaaggtccgagttgaagaacagcattgcggagtttcatacatatcaactcgatccggggagttgttctagtttgcacgcccaaa<br>ggatacacgccccaccggaacttgtgtggagtatcgtccgccggttcgataaaccacaaacacacaggcatttcattaaaagctgctccgtggagca<br>gaactttgaaatgagagtgggatgcacacgggatataatagttataagcggcctcccggccaacacgagcaccgaaagattggacatactggatga<br>cgagcgccgcgtgacgggggcatcaataatcggtggggaacatcgccttacgaattataaaggcgtcactactgttcatagatttgagaaggaaaat<br>agaatctggactgttgttctggaatcttatgtggttgatatgccagagggtaactcggaagatgatacacgcatgctggcggacacggttgtgaaacttaa<br>tctgcaaaagctggcaactgtcgccgaagccatggcaggtagcgcaggatcaggctcgagtggtaaagcatttatgcaagttgtcgaggctgacatg<br>ctctctaagggtctcctcggtggcgaggcgtggtcttcgtggcataaggaagactctattcatgtaggagtacgctgcatcgagatgctcattgagtcaac<br>cggaatggttagcctccaccgccaaaatgctggcgtagtaggtcaagactctgagactatcgaactcgcacctgaatacgctgaggctatcgcaaccc<br>gtgcaggtgcgctggctggcatctctccgatgttccaaccttgcgtagttcctcctaagccgtggactggcattactggtggtggctattgggctaacggtc<br>gtcgtcctctggcgctggtgcgtactcacagtaagaaagcactgatgcgctacgaagacgtttacatgcctgaggtgtacaaagcgattaacattgcgc<br>aaaacaccgcatggaaaatcaacaagaaagtcctcgcggtcgccaacgtaatcaccaagtggaagcattgtccggtcgaggacatccctgcgattg<br>agcgtgaagaactcccgatgaaaccggaagacatcgacatgaatcctgaggctctcaccgcgtggaaacgtgctgccgctgctgtgtaccgcaagg<br>acaaggctcgcaagtctcgccgtatcagccttgagttcatgcttgagcaagccaataagtttgctaaccataaggccatctggttcccttacaacatgga<br>ctggcgcggtcgtgtttacgctgtgtcaatgttcaacccgcaaggtaacgatatgaccaaaggactgcttacgctggcgaaaggtaaaccaatcggta<br>aggaaggttactactggctgaaaatccacggtgcaaactgtgcgggtgtcgataaggttccgttccctgagcgcatcaagttcattgaggaaaaccac<br>gagaacatcatggcttgcgctaagtctccactggagaacacttggtgggctgagcaagattctccgttctgcttccttgcgttctgctttgagtacgctgggg<br>tacagcaccacggcctgagctataactgctcccttccgctggcgtttgacgggtcttgctctggcatccagcacttctccgcgatgctccgagatgaggta<br>ggtggtcgcgcggttaacttgcttcctagtgaaaccgttcaggacatctacgggattgttgctaagaaagtcaacgagattctccaagcagacgcaatc<br>aatgggaccgataacgaagtagttaccgtgaccgatgagaacactggtgaaatctctgagaaagtcaagctgggcactaaggcactggctggtcaa<br>tggctggcttacggtgttactcgcagtgtgactaagcgttcagtcatgacgctggcttacgggtccaaagagttcggcttccgtcaacaagtgctggaag<br>ataccattcagccagctattgattccggcaagggtctgatgttcactcagccgaatcaggctgctggatacatggctaagctgatttgggaatctgtgagc<br>gtgacggtggtagctgcggttgaagcaatgaactggcttaagtctgctgctaagctgctggctgctgaggtcaaagataagaagactggagagattctt<br>cgcaagcgttgcgctgtgcattgggtaactcctgatggtttccctgtgtggcaggaatacaagaagcctattcagacgcgcttgaacctgatgttcctcggt<br>cagttccgcctccagcctaccattaacaccaacaaagatagcgagattgatgcacacaaacaggagtctggtatcgctcctaactttgtacacagcca<br>agacggtagccaccttcgtaagactgtagtgtgggcacacgagaagtacggaatcgaatcttttgcactgattcacgactccttcggtaccattccggct<br>gacgctgcgaacctgttcaaagcagtgcgcgaaactatggttgacacatatgagtcttgtgatgtactggctgatttctacgaccagttcgctgaccagtt<br>gcacgagtctcaattggacaaaatgccagcacttccggctaaaggtaacttgaacctccgtgacatcctcgagtcggacttcgcgttcgcgtaatagca<br>ccaccaccaccaccactgagatccggctgctaacaaagcccgaaaggaagctgagttggctgctgccaccgctgagcaataactagcataacccct<br>tggggcctctaaacgggtcttgaggggttttttgctgaaaggaggaactatatccg |
| pCDF-T7-<br>mcherry<br>(Turquoise:<br>T7 promoter<br>yellow: | agatctcgatcccgcgaaattaatacgactcactatagggagaccacaacggtttccctctacaaataattttgtttaactttaagaaggagatatacatat<br>ggtctccaagggcgaggaggacaacatggccatcatcaaggagttcatgcgcttcaaggtccacatggagggctccgtcaacgggcacgagttcga<br>gatcgagggcgagggggagggccggccgtacgagggcacccagaccgccaagctgaaggtgaccaagggcggccccctcccgttcgcctggg<br>acatcctctccccccagttcatgtacggctcgaaggcatacgtcaagcacccggccgacatcccggactacctgaagctctcgttcccggagggggttc<br>aagtgggagcgggtcatgaacttcgaggacggcggcgtcgtcaccgtcacccaggacagctccctgcaggacggcgagttcatctacaaggtcaa |

| | |
|---|---|
| mcherry<br>purple:<br>Terminator) | gctgcggggcacgaacttcccgagcgacggccccgtgatgcagaagaagacgatgggctgggaagcgtcctcggagcgcatgtacccggagga<br>cggcgccctcaagggcgagatcaagcagcgcctgaagctgaaggacggcggccactacgacgccgaagtcaagacgacgtacaaggccaag<br>aagccggtgcagctcccggGAgcctacaacgtgaacatcaagctcgacatcacctcgcacaacgaggactacacgatcgtggagcagtacgagc<br>gcgccgagggccggcactcgaccggcggcatggacgagctgtacaagtgagaattcttgttcagaacgctcggtcttgcacaccgggcgtttttctttg<br>tgagtcca |
